# Visualising nascent chain dynamics at the ribosome exit tunnel by cryo-electron microscopy

**DOI:** 10.1101/722611

**Authors:** Abid Javed, Tomasz Wlodarski, Anaïs. M.E. Cassaignau, Lisa D. Cabrita, John Christodoulou, Elena V. Orlova

## Abstract

Ribosomes maintain a healthy cellular proteome by synthesising proteins. The nascent chain (NC), emerges into the cellular milieu via the ribosomal exit tunnel, which is an active component that regulates the NC passage. How the NC dynamics at the exit tunnel affect NC folding remains to be an important question, the answer on which has strong implications to medicine. Here, we report high-resolution cryo-EM maps of ribosome nascent-chain complexes (RNCs) displaying distinct steps during biosynthesis. These RNC structures reveal a range of pathways adopted by the NC. The most pronounced diversity in the NC trajectories were found in the vestibule region. Rearrangements of several ribosomal components further suggest that these elements may actively monitor the emerging NC during translation. The ribosome-NC contacts within the vestibule define these NC pathways and modulate position of a folded immunoglobulin domain outside the ribosome.

## Introduction

Ribosomes are nano-machines that translate information coded in a messenger RNA into proteins in all living organisms. Recently, it has been found that ribosomes can also play a significant role in the process of co-translational folding by modulating the folding of a nascent chain (NC) during translation 1-6. Nascent chains (NC) can begin to acquire secondary structural elements in a co-translational manner during emergence via the ribosome exit tunnel within the large subunit of the ribosome 7, 8. Studies using cryo-EM have shown NC folding within the exit tunnel is largely limited to the formation of rudimentary secondary structure, predominantly α helices within different areas of the tunnel, while β hairpins as well as the formation of small domains in the wider vestibule region of the tunnel 9-13. Nonetheless, nascent polypeptides with more complex tertiary structure fold close to and outside the tunnel, as found for spectrin - a three-helix bundle protein, and titin, an all beta-sheet immunoglobulin domain 10,13. Biochemical and biophysical studies using FRET and PEGylation support these observations 14,15,16. Ribosome-nascent chain complexes (RNCs) studied by cryo-EM provided us with “snapshots” of most-stable states of NCs within the ribosomal tunnel 9-13. Yet, flexible features of the NC within the tunnel, especially at the ribosome tunnel vestibule, i.e. the lower tunnel, are poorly understood.

RNCs comprising the fifth and sixth domains of ABP-120 filamin protein (referred below as FLN5 and FLN6) have been analysed by NMR spectroscopy in the vicinity of the ribosome 17,18,19. In these RNCs, the FLN5 domain is tethered to the ribosome via different length sequences from FLN6 domain (Fig. 1a). Initial analyses of these RNCs revealed the transition of FLN5 to the folded state occurs when it is bound to the ribosome by a linker of approximately 45 amino acids 2, 20. Structural details on the organisation of FLN5 and FLN6 NC within the ribosome and the effect of the ribosome on the folding of FLN5 remains to be understood that would help to address the question on how the ribosome modulates co-translational protein folding.

**Figure 1.**
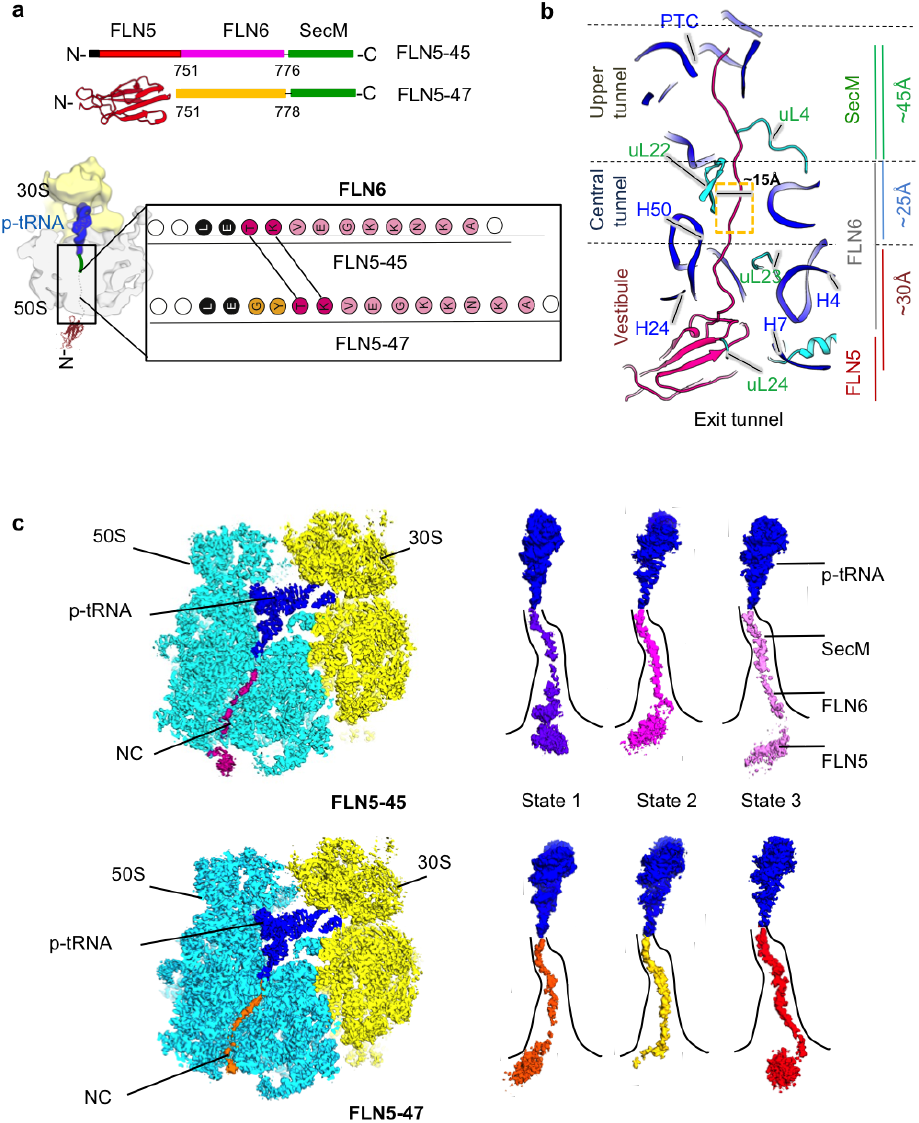
FLN5 RNC complex. a, Schematic diagram of RNC constructs with NC sequences: SecM (green), FLN6 (magenta/yellow) and FLN5 (PDB 1QFH, in red)23. Six histidines used for purification are shown in black, the two clonning residues between FLN6 and SecM sequence indicated with a line. The bottom right panel shows an organisation of the FLN5 NC at two linker lengths. The shift in FLN5-47 is due to two additional residues shown in yellow. b, Cross-section of the 50S subunit through the exit tunnel. Ribosomal RNA is in dark blue, ribosomal proteins are in cyan, FLN6 NC linker and FLN5 Ig domain are shown in magenta. The three main regions of the tunnel are indicated on the left. c, Upper left panel shows cryoEM map of FLN5-45 RNC (50S in cyan, 30S in yellow, p-tRNA in dark blue and NC in magenta). Upper right panel shows the EM density for three NC states of FLN5-45 within the ribosome exit tunnel: state 1 (purple), state 2 (magenta) and state 3 (pink). Bottom left panel shows the cryo-EM map of FLN5-47 RNC, NC in orange. Bottom right panel shows EM maps of three states of FLN5-47 RNC attached to p-tRNA; state 1 (orange), state 2 (yellow) and state 3 (red).

In this study, we report cryo-EM structures of SecM stalled FLN5-45 and FLN5-47 RNCs, where FLN5 is tethered to the ribosome by 45 and 47 amino acids using sequence from FLN6 and SecM. Statistical analysis of cryo-EM images allowed us to obtain a set of structures reflecting the most representative populations of RNCs at two points of translation. We have found that the NCs are dynamic within the tunnel and can adopt a variety of NC trajectories within the tunnel vestibule, resulting in distinct conformations for each NC linker within the exit tunnel, and three most probable positions of folded FLN5 NC. Trajectories adopted by the NC reveal the dynamic process of co-translational folding of the NC on the ribosome and in particular how interactions with the ribosome’s tunnel surface likely contribute to the productive folding of an emerging NC inside a cell.

## Results

### Ribosome-bound nascent chains adopt multiple trajectories

Two FLN5-RNCs were chosen as representative steps of FLN5 NC (residues 646 to 750) biosynthesis, which is tethered to the ribosome by 45 or 47 amino acid linkers that comprise either 26 (A751 to T776) or 28 (A751 to G778) residues of FLN6 together with the 17 amino acid SecM stalling sequence 21 (Fig. 1a). Cryo-EM images for FLN5-45 and FLN5-47 RNCs samples were collected on a Titan Krios electron microscope (Thermo Fisher ScientificTM), operating at 300 kV and equipped with a K2 Summit direct electron detector (see Methods; Table 3). We used RELION 2.0 for structural analysis of the RNCs (see Methods, Supplementary Fig. 1). The local resolution obtained from the maps of FLN5-45 and FLN5-47 RNCs displayed both stable and flexible regions in both the ribosome and NC (Supplementary Fig. 2). In the structures obtained, the two ribosomal subunits have an average resolution of ~ 3.5 Å. The resolution of NC within the tunnel was lower compared to the overall resolution of the ribosome, suggesting that FLN6 region of the NC is highly dynamic (Supplementary Fig. 2). Further 3D classification of RNCs enabled us to distinguish structures according to the location of the FLN6 domain within the vestibule region of the ribosomal exit tunnel (Fig. 1; Supplementary Fig. 1, see Methods,).

Since the RNC complexes analysed here were in the SecM-arrested state, it enabled us to characterise features of the NC within the tunnel. The distinct density corresponding to the NC spans the entire ~ 100 Å length of the exit tunnel. The tunnel can be divided into three major regions depending on the distance from PTC (Fig. 1b): upper - between the entrance and the tunnel constriction formed by uL4 and uL22; central - between the tunnel’s constriction site and nucleotide A1321 of H50 (starting at ~ 45 Å from the tunnel entrance); and the vestibule region (starting at ~ 70 Å from the top, and ~ 25 Å wide) (Fig. 1b). The P-site tRNA, located between the small and large ribosomal subunits in the peptidyl transferase centre (PTC), is well defined, as it is covalently linked to the NC at the entrance to the exit tunnel (Fig. 1b). The structures show the P-site tRNA (in blue, Fig. 1c) is attached to the NC via the Gly165 residue, which is the penultimate residue observed in the SecM sequence at the entrance to the tunnel. Density corresponding to the SecM sequence of the NC threads throughout the upper tunnel region (~ 10 Å wide). This NC density could be traced further within the central tunnel part and was assigned to the FLN6 sequence. In this region of the tunnel, the resolution was sufficient in both RNCs to clearly model side chains of residues 776-773 (FLN5-45) and 778-774 (FLN5-47) in FLN6 (Fig. 2). Although the NC densities inside the wider vestibule area (~ 25 Å) became more diffused, we were able to trace the remaining 22 (FLN5-45) and 24 (FLN5-47) residues of FLN6 as a poly-alanine backbone. Analysis of the densities corresponding to the NC (based on the classification of EM data in RELION) 22 enabled us to define three trajectories adopted by the NC within the vestibule region (Table 1; Fig. 1). A bulk of density in both FLN5-45 and FLN5-47 RNCs linked to FLN6 and observed close to the tunnel’s exit and the ribosome surface was attributed to FLN5 NC. Subsequent fitting of the FLN5 domain crystal structure 23 into this outer density indicates that the volume of these densities is consistent with folded FLN5 (Table 2; Fig. 1c).

**Figure 2.**
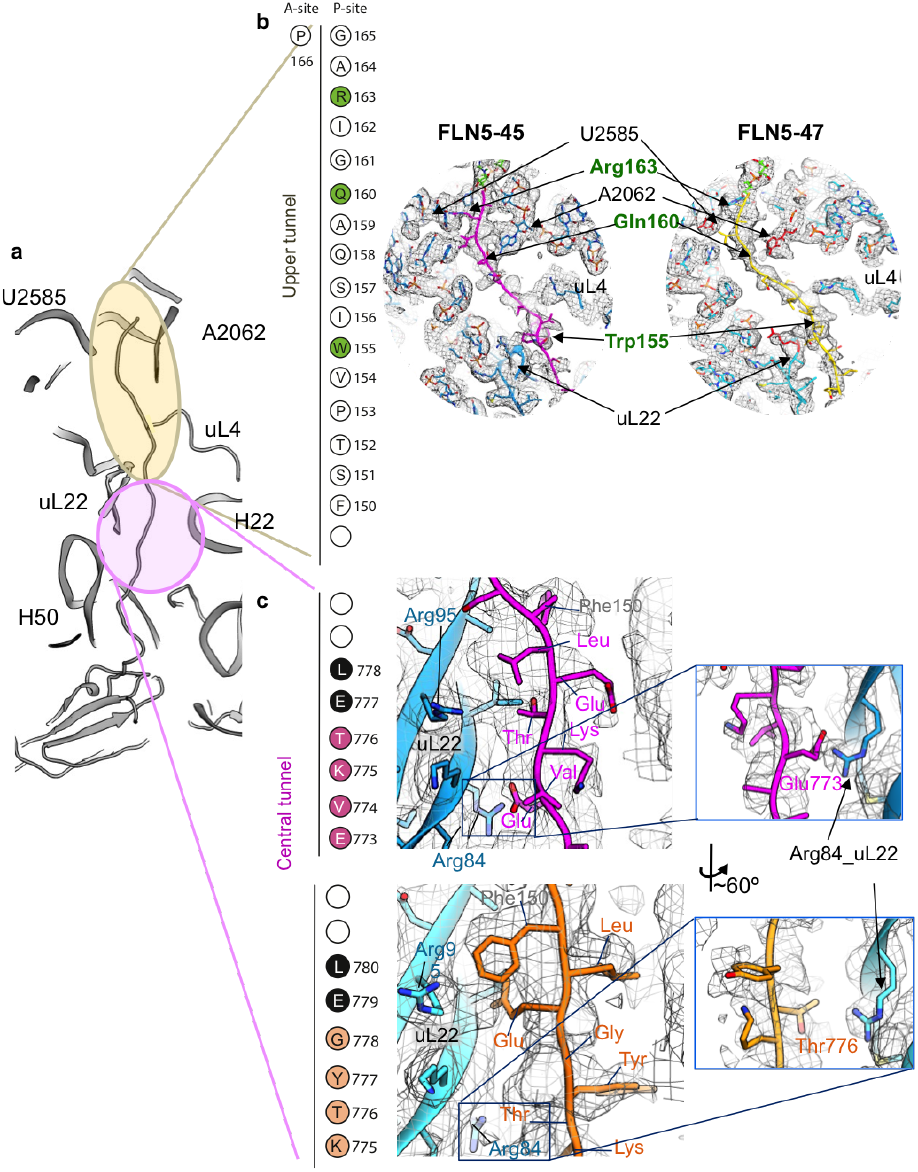
SecM and FLN6 NC within the tunnel.a, Cross-section of the exit tunnel. Regions corresponding to upper and central tunnel are shown in yellow and pink respectively. b, Left panel shows 17 residues of the SecM NC. C-terminal proline is located at the ribosome A-site, linked to Gly-stalled SecM NC. Key interacting residues are coloured in green. Right panels show the upper tunnel regions from FLN5-45 (state 2, in magenta) and FLN5-47 (state 2, yellow) RNCs. Ribosomal RNA and proteins are coloured in light blue and NC in magenta (FLN5-45) and yellow (FLN5-47). Nucleotides and protein residues that interact with SecM NC are shown in red and labelled in green. c, Close-up cross sections of central tunnel in FLN5-45 (state 2 – magenta) and FLN5-47 (state 2 – yellow). Residues of uL22 (blue) and NC are labelled. Middle panel shows the residues of FLN6 fitted within the map in FLN5-45 (magenta) and FLN5-47 (yellow). Right panel shows close-up views of interactions between FLN6 NC residues and uL22-Arg84 (blue).

**Table 1.**
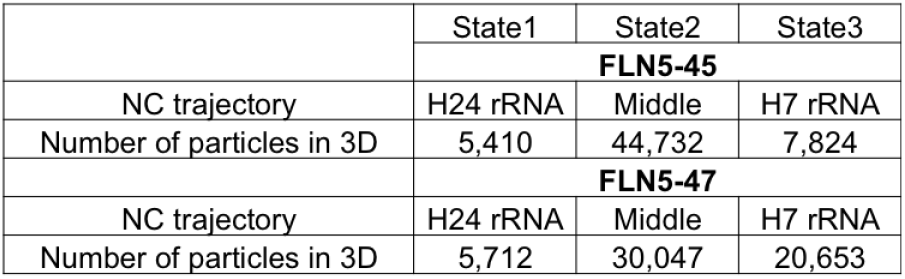
NC states identified in FLN5 RNC complexes.

|  | State1 | State2 | State3 |
| --- | --- | --- | --- |
|  | **FLN5-45** |  |  |
| NC trajectory | H24 rRNA | Middle | H7 rRNA |
| Number of particles in 3D | 5,410 | 44,732 | 7,824 |
|  | **FLN5-47** |  |  |
| NC trajectory | H24 rRNA | Middle | H7 rRNA |
| Number of particles in 3D | 5,712 | 30,047 | 20,653 |

**Table 2.** Fitting scores for FLN5 in EM maps of FLN5-45 and FLN5-47 RNCs.

|  | State1 | State2 | State3 |
| --- | --- | --- | --- |
|  | **FLN5-45** |  |  |
| CC-fit Model to EM map | 0.72 | 0.78 | 55 |
|  | **FLN5-47** |  |  |
| CC-fit Model to EM map | 0.75 | 0.45 | 0.76 |

Based on the identified NC trajectories within the vestibule three NC pathways that were identified are referred further as “states” (1 to 3) for each RNC length. State 1 corresponds to a population of RNCs in which FLN6 is in close proximity to H24 of the 23S rRNA in the vestibule, state 2 represents a NC that is positioned between H24 and H7 23S rRNA, while state 3 corresponds to FLN6 NC located close to H7 23S rRNA; the relative populations of the observed FLN5-45 and FLN5-47 states in the cryo-EM datasets are shown in Table 1. Each of the states observed is accompanied by small local changes in the ribosomal elements lining the vestibule (see below). Analysis of both complexes shows that states 2 are the most populated subsets (Table 1), which likely represent the most long-living and stable conformation of the NC for these two RNCs complexes.

### Dynamic properties of the SecM stall motif

All structures of the FLN5-RNCs show well-defined density within the upper and central tunnel regions occupied by the SecM NC (Gly165 - Phe150) (Fig. 2a, b). The initial fit was done using NC states 2, where SecM was resolved on average at ~ 5 Å. These models were subsequently used to trace SecM in less populated states in both RNCs. These fittings of the SecM sequence into the densities of the NC revealed stable interactions with the ribosomal exit tunnel that induce translational arrest, as established by previous studies 21, 24, 25 (Fig. 2b; Supplementary Fig. 3). The interaction between Arg163 of SecM and nucleotides U2506 and A2451 (Int163,2506) of 23S rRNA at the PTC in both FLN5-45 and FLN5-47 RNC complexes was well defined (Fig. 2b; Supplementary Fig. 3bi, di). Close to the PTC in the upper tunnel, SecM Gln160 appears to interact with A2062 in FLN5-45 RNC, not observed in FLN47-RNC, highlighting flexibility of SecM and variable, non-specific interactions within ribosome tunnel (Fig. 2b; Supplementary Fig. 3bii, dii). Another interaction is found between Trp155 (SecM) and A751 (23S rRNA) (Int155,751), located close to the uL4-uL22 constriction site, that is visible in both structures of FLN5-45 and FLN5-47 RNC (Fig. 2b; Supplementary Fig. 3biii, diii). This NC density is in a good agreement with reported structural and mutagenesis studies indicating that Trp155 to Ala155 mutation reduces SecM stalling 26, 24, 25. The distances between Int163,2506 and Int155,751 of SecM (20-22Å), were found to be shorter than the expected ~ 30 Å for an extended SecM NC chain of the same length (assuming that the distance between two residues in an extended chain is ~ 3.4 Å) (Supplementary Fig. 4a). This indicates that SecM NC in the FLN5 RNCs is in a compact form, which is consistent with previous studies suggesting this effect to exist for NC within the ribosome exit tunnel (Supplementary Fig. 4a) 27, 28, 25.

We further found structural differences in the SecM NC conformations of the two RNCs. This dynamic behaviour of SecM is seen at the tunnel constriction (uL4/uL22 constriction site); we found that the N terminal residues (Pro153–Ser151) of the SecM chain undergo lateral shifts of ~ 3.8 Å between states 2 of FLN5-45 and FLN5-47 RNCs (Supplementary Fig. 4b). This shift of the polypeptide chain is possibly related to the residue Pro153 of SecM, the mutation of which makes SecM more flexible within the tunnel, supporting biochemical data on its role in mediating translational arrest NC 29. Observing this variability in SecM NC position nearby the tunnel constriction site is also in agreement with previously reported molecular dynamics simulation of SecM NC within the tunnel 28. It suggests that the shift of Pro153 observed in our SecM NC structures is possibly arising from the NC flexibility downstream in the vestibule. Altogether, the key interactions identified in our structures confirm previously described structures of SecM-arrested RNCs 24, 25 indicating that SecM is essential for the translational stalling, but further show that SecM NC can be affected by flexibility of linked NC downstream within the tunnel.

### FLN6 NC contacts with the central tunnel

The central tunnel region located beyond the tunnel constriction site is slightly wider and has a diameter of ~ 15 Å. It is lined with nucleotides of 23S rRNA domains H22, H23, H50 and uL22, uL23 ribosomal protein segments (Fig. 1b). The structures of the FLN5-45 and FLN5-47 RNCs show NC density along the central tunnel region at a resolution of ~ 6.0 Å (Fig. 2c; Supplementary Fig. 2) allowing us to trace the backbone of the FLN6 polypeptide chain. All states of FLN6 NC in both RNC lengths show that EM densities of the NC are, in general, similar but vary in local interactions with the central tunnel elements. In the RNCs, this tunnel part accommodates about 10 residues of FLN6 NC: two residues that link the SecM and eight residues of the C-terminal FLN6 NC. We have fitted atomic models of the two linker residues (Leu-773 and Glu-774) and four out of eight FLN6 residues (Thr776-Glu-778 in FLN5-45 and residues Tyr-777 to Lys-778 in FLN5-47; see methods, Fig. 2c), the remaining residues were modelled with poly-alanine. Fittings of FLN6 NC within the central tunnel were derived from the most populated states, state 2 in both in both FLN5-45 and FLN5-47 RNCs.

Structures of FLN5 RNC state 2 show that FLN6 NC makes contacts with the central tunnel via the ribosomal protein uL22 (Fig. 2c). While in the FLN5-45 RNC, Glu-778 in FLN6 appears to interact with Arg84 of uL22 (Fig. 2c, upper panel), in FLN5-47 it is Thr-776 of FLN6 NC that interacts with uL22 (Arg84) (Fig. 2c, bottom panel). Known structures of RNCs with different sequences of NCs also show interactions with Arg84 of uL22 11,12, indicating that uL22 acts as a non-discriminatory point of interaction between the ribosome tunnel and the NC.

### Interactions of FLN6 NC at the tunnel vestibule entrance

At the entrance to the tunnel vestibule, lined by A1321 nucleotide of H50 23S rRNA and the beta-hairpin loop of uL23 reveals changes between FLN5-45 and FLN5-47 RNCs in contact points made by FLN6 NC (Fig. 1b, Supplementary Fig. 5,). These were most pronounced in NC states 2 of FLN5-45 and FLN5-47 RNC and state 3 of FLN5-47 RNC (Supplementary Fig. 5). In state 2 of FLN5-45 RNC, the FLN6 NC interacts with both A1321 nucleotide and uL23 (Supplementary Fig. 5a), but in FLN5-47 RNC (state2), only one contact is observed with A1321 (Supplementary Fig. 5b). Interestingly, state 3 of FLN5-47 RNC shows two contacts of NC with H50 and uL23, similar to state 2 FLN6 of FLN5-45 RNC (Supplementary Fig. 5c). It is likely that several positively charged residues in FLN6 NC sequence located at the vestibule (i.e. Lys771-Lys770-Lys768) make some electrostatic interactions with negatively charged 23S rRNA (Fig. 3a; Supplementary Fig. 5). Difference in NC pathways at the tunnel vestibule have been previously observed for stalled NC with different sequences 30, suggesting that the vestibule appears to be a non-discriminatory region that may modulate the exit route of the translated NC. Apparently, the strength of the uL23-H50 rRNA interaction with the NC plays a role in the trajectory shaping of the NC at the vestibule, shifting it either towards H7 (state 3) or exiting via the middle of the tunnel (state 2).

**Figure 3.**
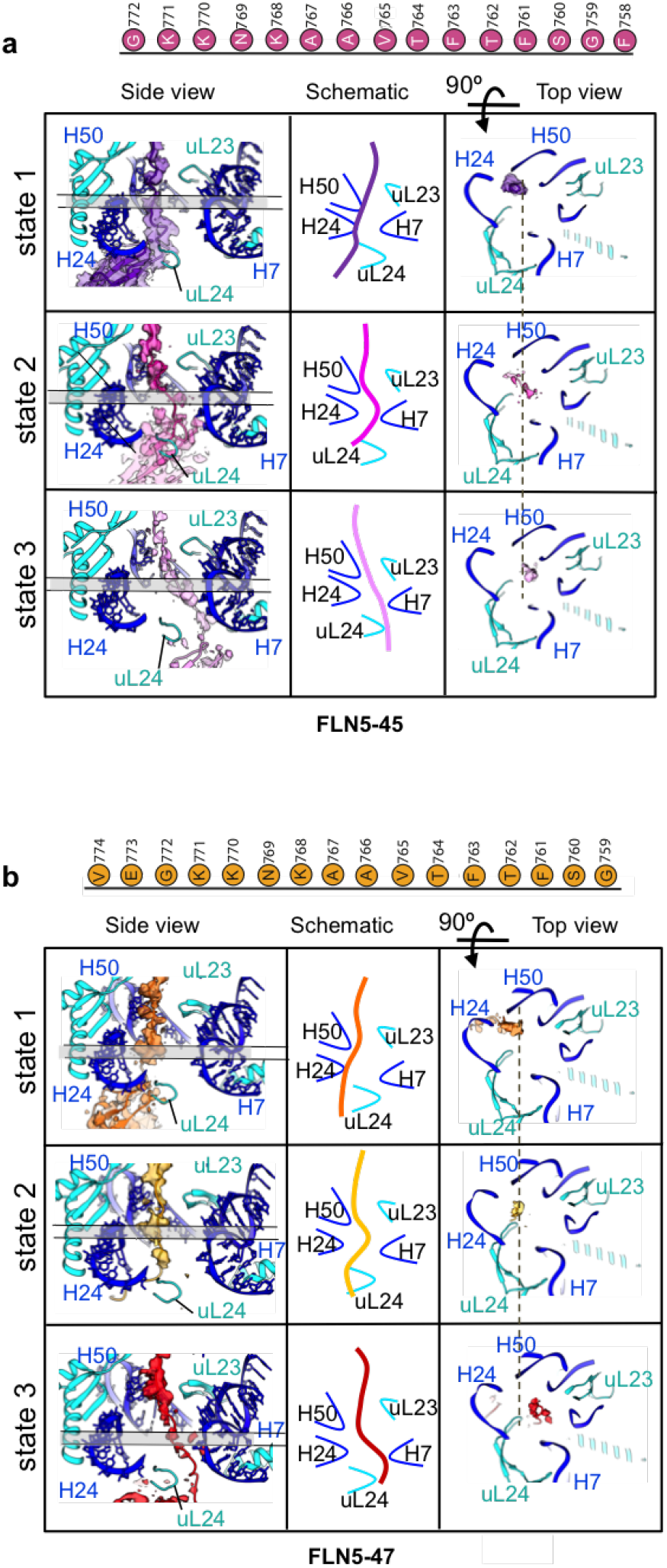
NC trajectories within the vestibule. a. States 1-3 is for FLN5-45. Left panels show the cross-section along the ribosome exit tunnel at the area of the vestibule, middle panels show schemes of interactions between NCs and ribosomal elements lining the tunnel; right panels show the cross section of the NCs within the tunnel on the level of the shaded areas shown in the left panels correspondingly. The dashed line is centered according to NC states 2 and indicates shifts in positions of NC in both a and b. b. States 1-3 is for FLN5-47. The same representation of the NCs within the exit tunnel as in a. Ribosomal proteins are shown in cyan, the NCs are colour coded as in Figs. 1 and 2. Sequences corresponding to FLN6 in the vestibule in FLN5-45 (in magenta) and in FLN5-47 (in yellow) are shown on top panels in a, and b, respectively.

### NC trajectories within the tunnel vestibule

Across the different RNC states, a point of divergence of FLN6 NC trajectories was found at A1321 nucleotide of H50 (23S rRNA) (Fig. 3). These non-specific, charge-mediated interactions appear to produce multiple NC positions within the tunnel vestibule. Three distinct and most long-living states in both RNCs were identified: one is in close proximity to the H24 rRNA domain (state 1), the second at the centre of the vestibule (state 2) and the last one being close to the H7 rRNA domain (state 3, Fig. 3, Supplementary Fig. 6b). The ribosome tunnel vestibule is lined by negatively charged 23S rRNA (nucleotides of H50, H24, H7 and H4) and loops with positively charged residues from uL23, uL24 and uL29 proteins (Fig. 1b). The large diameter of the vestibule permits greater flexibility to the NC, as observed in the different RNC states (Fig. 1; Table 1). The resolution of FLN6 in our FLN5-RNCs in the vestibule was of ~ 8.0 Å, which permitted to trace the N-terminal portion of FLN6 NC as a poly-alanine chain (from residue Lys771 onwards towards the C-terminal Ala751) (Fig. 3).

Examination of the trajectories of FLN6 NC within the vestibule has shown that the N-terminal region (from residues Ala767 onwards) of FLN6 NC in states 1 in both FLN5-45 and FLN5-47 RNCs interacts with nucleotides of H50 and H24 domains of 23S rRNA and uL24 loop (Fig. 3). Interestingly, FLN5-45 NC state 1 has a bulk of density attached to nucleotides H50-H24 of 23S rRNA, which is likely, related to a compaction of FLN6 NC (Fig. 3; Supplementary Fig. 6a). This state 1 is possibly short-lived and, therefore, the least-populated NC state. State 3 of FLN6 NC in both FLN5-45 and FLN5-47 RNCs shows several contacts with uL23 and uL29 loop and with the nucleotides of H4 and H7 domains of 23S rRNA (Fig. 3).

Positions of FLN6 NC varied significantly in the vestibule and to assess the shifts in different states, the states 2 NC (the most populated in both RNCs) were used as the reference point (Table 1, Supplementary Fig. 6b,c). In FLN5-45 RNC, state 1 NC is shifted by ~ 7 Å (from the centre of the tunnel to the H24 side) and state 3 is shifted by ~ 8 Å (from the centre to the H7 side) with respect to state 2. A similar effect is observed in FLN5-47 in which states 1 (towards H24) and 3 (towards H7) are shifted by 5 Å and 7 Å, respectively, relative to state 2 (Supplementary Fig. 6b,c). Overall, the shifts of the NC within the vestibule in both FLN5-45 and FLN5-47 RNCs indicate large movements (up to ~ 15 Å), that are consistent with the 20 Å wide vestibule (Supplementary Fig. 6b,c). The multiple NC trajectories at the vestibule suggest that the translated NC is flexible, and has multiple contact points within the tunnel that forms temporary conformations of FLN6 NC at the vestibule. As a result, the movement of FLN6 NC within the vestibule directly affects positions of linked FLN5 NC located close to the tunnel exit (as described below).

### The ribosomal tunnel vestibule is active

Our RNC structures reveal variability in positions of ribosome protein loops uL23, uL24 and uL29 across different RNC states. Comparison of states 2 of FLN5-45 and FLN5-47 RNCs shows that the uL23 loop moves by ~ 3 Å (Fig. 4a). Interestingly, the uL24 tunnel loop, located at the mouth of the tunnel exit position, in the RNC states undergoes shifts from 3 to 5 Å. Between states 2 of FLN5-45 and FLN5-47 RNCs the uL24 loop is shifted by ~ 5 Å (Fig. 4b). This shift indicates that the uL24 loop is flexible, which is consistent with recent RNC structures reporting similar shifts in uL24 loop in the presence of NC (Tian et al., 2018). The loop of uL29 has the smallest shift (~ 1 Å) between states 2 of FLN5-45 and FLN5-47 (Fig. 4c). A biochemical study has shown that the uL23, uL24 and uL29 loops can be cross-linked with emerging NCs, indicating that they interact with an emerging NC 31. In addition to the changes in positions of loops from uL23, uL29 and uL24, small shifts (~ 1 to 2 Å) were also detected in several regions in the 23S rRNA, including A1321 of H50, A91 nucleotide of H7 in 23S rRNA and A490 nucleotide of H24 (Supplementary Fig. 7).

The comparison of our FLN5 RNC structures to an empty 70S ribosome (Noeske et al., 2015) has shown that the ribosomal protein loops in RNCs undergo substantial shifts: uL23 moves by ~ 12 Å (from the empty 70S to the RNC) and uL24 by ~ 6 Å (Fig. 4a). Similarly, the local shifts are also observed in the nucleotides of rRNA domains that interact with FLN6 NC at the vestibule: H24 (A490) and H50 (A1321) move by ~ 3 Å, and H7 (A91) by ~ 2 Å (Supplementary Fig. 7). The changes in these rRNA regions indicate that the NC induces local rearrangements due to non-specific interactions with the ribosome tunnel.

**Figure 4.**
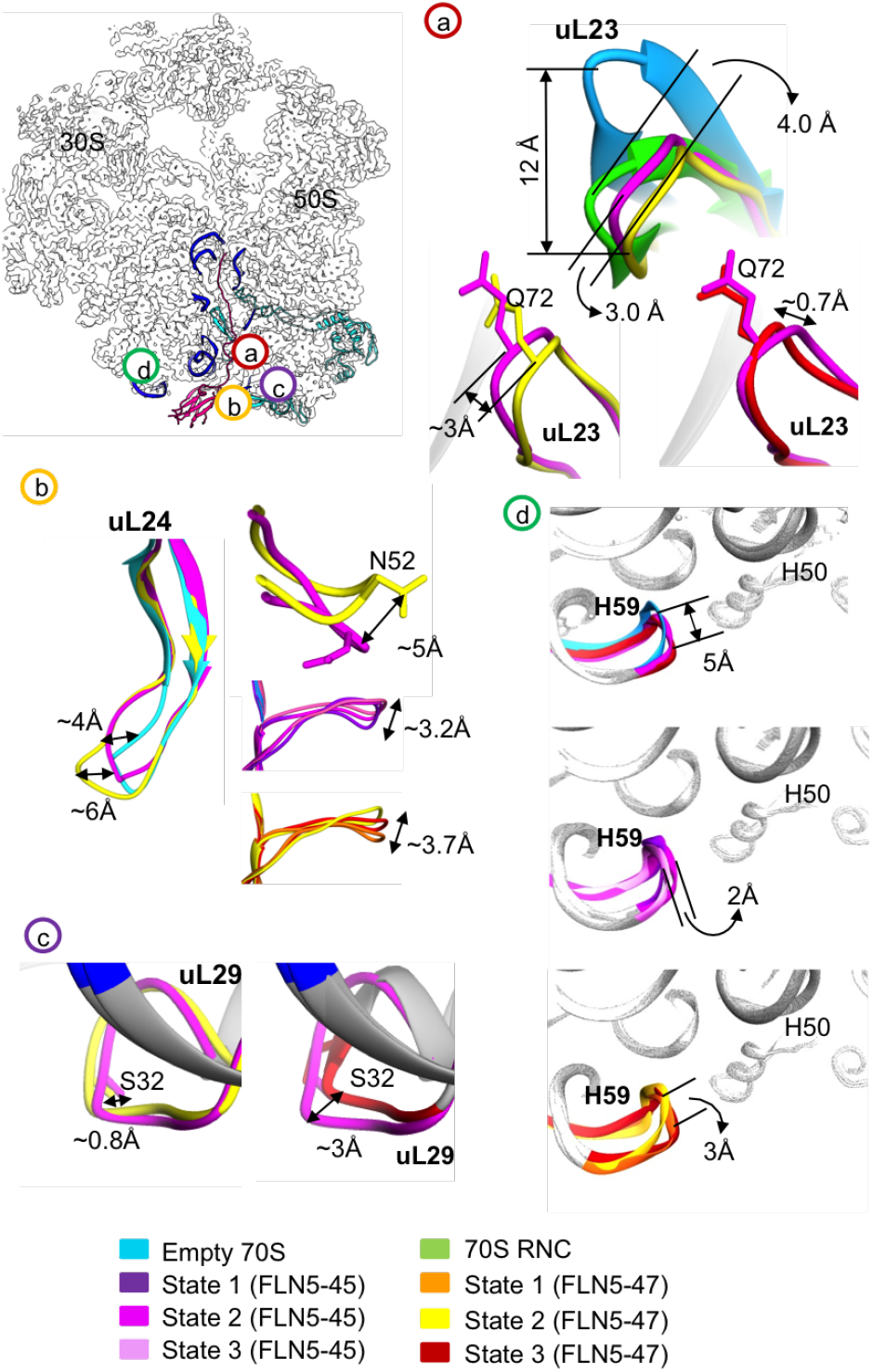
Changes in ribosome tunnel vestibule in FLN5 RNCs. Top right panel shows the ribosome map in grey, labelled with elements of the exit tunnel (RNA in blue and proteins in cyan). Structural changes in the exit tunnel are shown in panels a, b, c and d. a, changes in uL23; b, changes in uL29; c, changes in uL24; d, changes in ribosomal RNA domain H59.

### FLN5 NC is folded in the vicinity of the ribosome exit tunnel

The cryo-EM structures of FLN5-FLN6 RNCs show a distinct bulk of density just outside the vestibule. We proposed this density to be of folded FLN5 domain and verified it by rigid-body fitting of the X-ray structure of FLN5 domain (PDB 1QFH; Fig. 5a, Supplementary Fig. 8a; see methods). Assessment of the cross-correlation (table 2) between the X-ray crystal structure and the EM density shows that the locations and sizes of these bulks of densities are consistent with the immunoglobulin domain and therefore it was assigned to the FLN5 domain (Fig. 5a).

**Figure 5.**
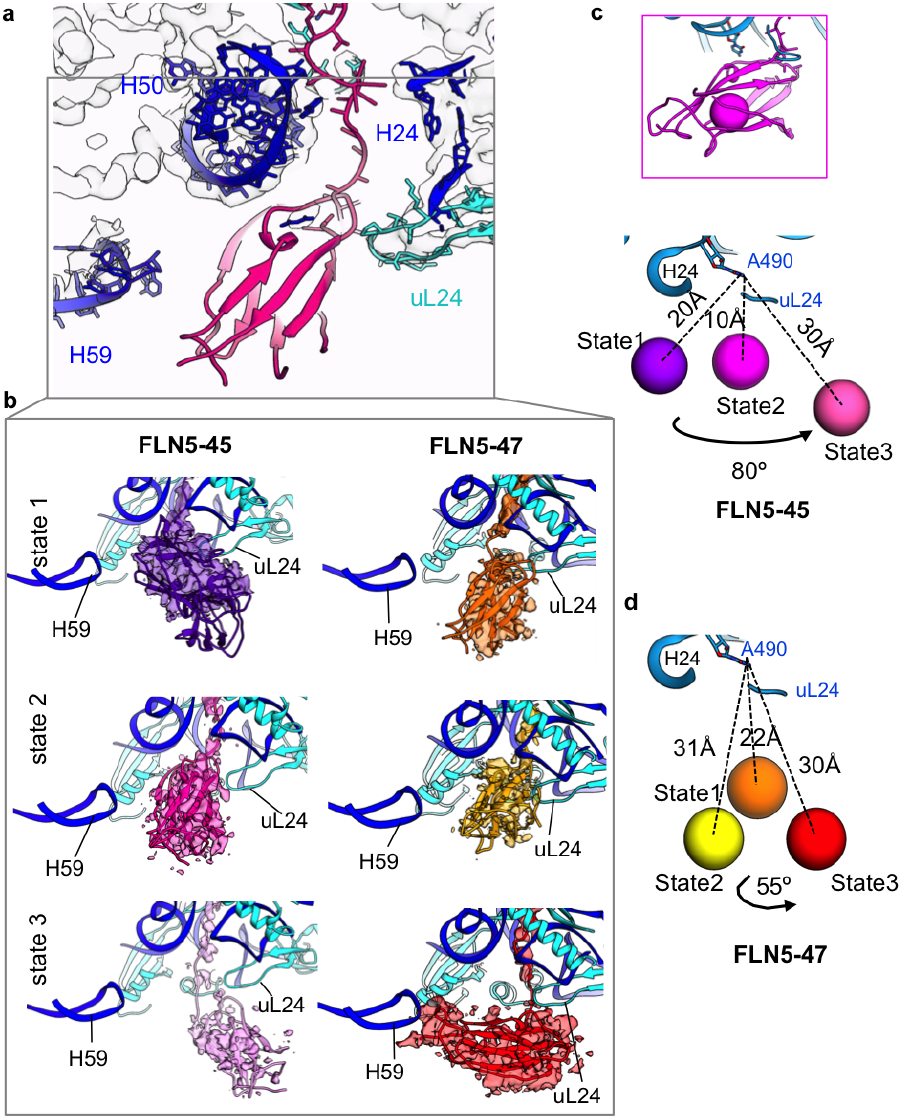
Changes in positions of FLN5 domain on the ribosome. a, A close up view of the tunnel vestibule with modelled FLN5 domain (FLN5-45 state 2). b, FLN5 domain positions for FLN5-45 states (left column) and for FLN5-47 states (right column). c, Models of FLN5 in three states (shown as spheres located at central mass of the FLN5 domains, as depicted in the upper panel) for FLN5-45 with distances from the tunnel exit and the FLN5 centres and the degree of rotation of FLN5. d, Schematic representation of FLN5 shifts in the FLN5-47 RNCs between three states. Measured distances of FLN5 position from the tunnel (A490 nucleotide on H24 rRNA domain) are indicated. Ribosome elements are shown in a colour scheme as in the previous figures.

The structures of RNCs in different states revealed variations in the position of the FLN5 NC near the ribosomal exit tunnel (Fig. 5). In states 1 and 2, FLN5 domain is positioned at ~ 16 to 20 Å away from H24 ribosome domain, and close to the uL24 ribosome tunnel loop (Fig. 5b). By contrast, FLN5 in state 3 of FLN5-45 RNC is positioned ~ 32 Å away from the ribosomal surface (using H24 as a reference point), and located close to the uL29 protein on the ribosome surface (state 3, Fig. 5b). In FLN5-47 RNC states, the FLN5 domain in state 1 is ~ 22 Å away from the tunnel exit and in the proximity of H24 rRNA domain (Fig. 5b). We note that the point of attachment of FLN5 in state 1 of FLN5-47 to the ribosome at H24 rRNA domain is analogous to that of FLN5 observed in states 1 and 2 of FLN5-45 RNC (Fig. 5b). The less defined densities of FLN5 in state 2 of FLN5-47 indicate that FLN5 NC is shifted by ~ 15 Å away from the ribosomal surface and is dynamic (Fig. 5c). In state 3 of FLN5-47 RNC, FLN5 is located ~ 30 Å away from H24 of the ribosome tunnel (Fig. 5c). The FLN5 NC in this state is rotated by ~ 60 degrees clock-wise towards H59 rRNA domain relative to the FLN5 domain in state 2 of FLN5-45 RNC (state 3, Fig. 5a, bottom panel). FLN5 in state 3 of FLN5-47 RNC also shows that the FLN5 N-terminus interacts with H59 of the 23S rRNA domain (Fig. 5a, bottom panel; Supplementary Fig. 8b). The contact is possibly mediated by electrostatic interactions of the negatively charged H59 rRNA domain nucleotides and positive charges (corresponding to Lys and Arg residues) in the N-terminal loops of FLN5 (Supplementary Fig. 8b). We found that this FLN5 NC interaction with the H59 domain induces a shift of ~ 5 Å in the rRNA loop, compared to H59 loop in the empty ribosome, and a shift of ~ 3 Å in H59 rRNA loop compared with state 2 of FLN5-47 RNC structure (Fig. 4d). This contact of NC with H59 rRNA domain has also been observed in a partially folded NC system (Nilsson et al., 2017). Henceforth, interaction of NCs with H59 rRNA domain suggests H59 plays a role in co-translational events as the NC emerges from the exit tunnel during translation.

The states observed for this tandem FLN5-FLN6 RNC are defined by non-specific interactions of the FLN6 domain with the ribosome tunnel, which modulate positions of the FLN5 domain. The shifts in positions of FLN5 NC in the vicinity of the exit tunnel for both FLN5-45 and FLN5-47 RNCs correlate with the FLN6 NC trajectory within the vestibule and reflect the NC dynamics linked to translation (Fig. 3; Supplementary Fig. 6).

## Discussion

The cryo-EM structures of RNCs obtained in this study comprise a tandem of two immunoglobulin domains FLN5 and FLN6 (the full length of FLN5 (105 residues) and 26 and 28 residues from FLN6) from ABP120 protein (see methods) attached to the stalled ribosome by SecM NC. Densities of the NCs within the ribosomal exit tunnel are traced and demonstrate for the first time the dynamics of the nascent polypeptide chain along the entire length of the tunnel. These structures show three most populated trajectories (states) in each RNCs (i.e. FLN5-45 and FLN5-47).

The two additional residues in the FLN6 (FLN5-47 RNC) tether shift the folded FLN5 NC domain slightly further away from the ribosome, enhancing its ability to change its position nearby the exit tunnel and facilitate temporary points of interaction with a peripheral ribosome region (i.e. the H59 domain of 23S rRNA). Comparison of the different states of two RNCs shows that the FLN6 tether appears to be in a compact state in the vestibule and due to its flexibility, FLN6 affects the position of FLN5 around the tunnel exit. Positions occupied by FLN5 NC also are found to be close to the ribosome exit tunnel on average 10-30 Å from the ribosome surface compared with the expected 60-70 Å distance from the tunnel exit for the extended tether. The compaction of the FLN6 and the close position of FLN5 to the ribosome surface may explain difficulties in observations by NMR of the FLN5 folding in complexes with the tether of 21-42 aa compared to the folded polypeptide in solution 2. Possibly, FLN6 NC compaction and non-specific interactions of the NC inside the exit tunnel (observed in our structures) contributes to an impeding premature folding and as a result sequestering part of the FLN5 NC sequence inside the tunnel at short tethers.

Analyses of the RNC structures show three sites along the ribosome exit tunnel that may mediate NC activity within and close to the exit tunnel (Fig. 6). These include uL22 at central tunnel, uL23-H50-H24 at the vestibule and H59 outside the tunnel. An electrostatic interaction between H59 and FLN5 in FLN5-47 RNC apparently stabilises one of the positions of FLN5 NC. Published cryo-EM structures of RNCs also implicate uL22 and H59 as main interaction points for an emerging NC, influencing the NC position within the tunnel and affect NC folding 10,11,12. Interesting parallels can be drawn with recent cryo-EM studies of RNCs of small folded domains on shorter linker lengths: both spectrin (linker length 32aa) 10 and the immunoglobulin-like protein, I27, (linker length 35aa, Tian et al, 2018). Notably, spectrin and I27 domains show a preference for the orientation found in FLN5-47 state 3, suggesting that specific sites of contact on the ribosome of the newly formed globular state exist to favour co-translational folding.

The multitude of NC states on the ribosome confirms the fact that NC is dynamic in the vestibule and its conformational flexibility is increased. With enlarged diameter of the tunnel towards its exit in bacterial ribosomes, the likelihood of short-lived, non-specific interactions between the ribosome tunnel and the NC increases.

**Figure 6.**
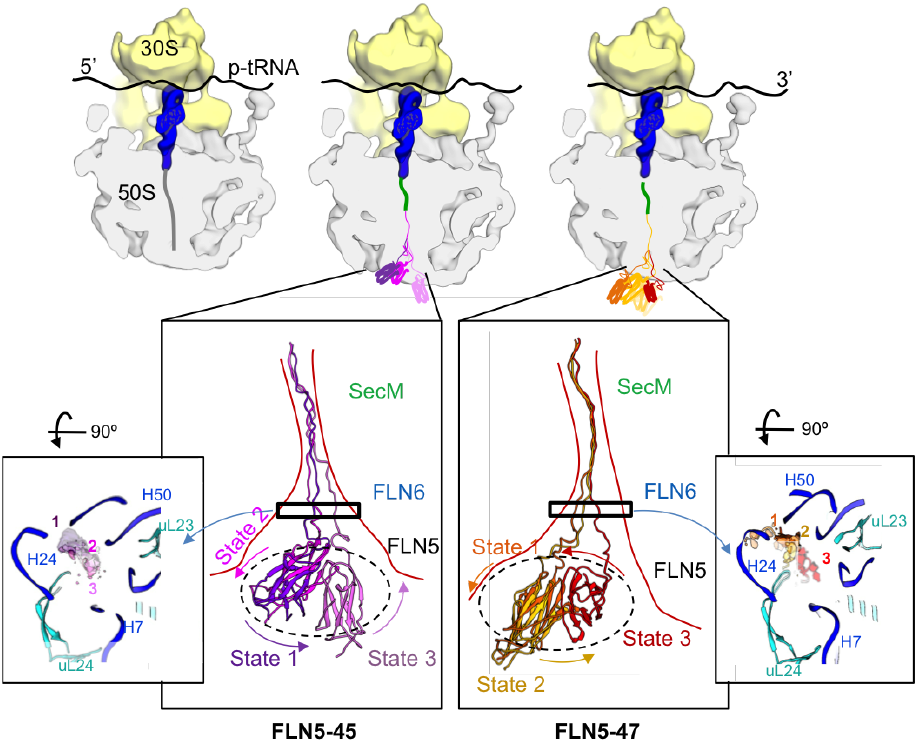
Schematic representation of the NC mobility within the exit tunnel. Upper panel shows a schematic diagram of the ribosomes during translation of FLN5 RNC constructs in three steps; step one translating SecM, step two translating FLN5-45 RNC (with its three main states in the zoom-in panel) and step three translating FLN5-47 RNC (three states in the zoom-in panel). Bottom panel shows two zoom-in views into both FLN5-45 and FLN5-47 RNC NC states: first panel shows the FLN6 trajectories inside the tunnel vestibule and second panel for each states shows the rotational movement of FLN5 around the tunnel exit.

## Ackowledgements

This work was supported by the Wellcome Trust (206409/Z/17/Z) to JC; Dr A. Javed by MRC DTP PhD grant MR/J003867/1. We would like to thank Dr Dan Clare and Dr Alistair Seibert at eBIC (Diamond) who assisted in cryo-EM data collection. We thank Dr D. Houldershaw, R. Westlake, and Y. Goudetsidis for computer support throughout the duration of the project. We would also like to thank Prof G. Waksman for productive discussions of the project.

## Author Affiliation

E.V.O., L.D.C., and J.C. designed the research. E.V.O. and J.C. supervised the overall project. L.D.C. and A.M.E.C. purified the RNC constructs. A.J. prepared cryo samples, collected data, processed and did structural analysis. A.J. and T.W. have done interpretation of structures obtained. A.J., L.D.C, J.C. and E.V.O wrote the manuscript. All authors discussed the results and contributed to the final version.

Three-dimensional cryo-EM density maps of the RNCs complexes have been deposited in the Electron Microscopy Data Bank under the accession numbers EMD-XXXX, EMD-XXXX, EMD-XXXX, EMD-XXXX, EMD-XXXX, EMD-XXXX. Atomic coordinates of the RNCs for each state of the complexes have been deposited in the RCSB Protein Data Bank under the accession codes YYYY, YYYY, YYYY, YYYY, YYYY and YYYY. Correspondence and requests for materials should be addressed to E.V.O. or J.C.

## Materials and Methods

### Generation of ribosome-nascent chain complexes (RNCs)

RNC constructs of FLN5-45 and FLN5-47 for were derived from a FLN5-110 RNC construct by truncating the C-terminal residues in FLN6 linker sequence, as described previously 2. The constructs include 17 residues from SecM, two cloning residues, 26/28 residues from FLN6 and the full 105 sequence of FLN5 from ABP120 protein 2. The FLN5-RNCs were biochemically purified as described in 20, though with included minor modifications: following growth in an un-labelled MDG medium at 37°C, the cells were washed and re-suspended in EM9 medium. RNC expression was induced with 1 mM IPTG at 30°C, and 150 mg/mL rifampicin was added 10 minutes later. The cells were harvested by centrifugation 35 min later. Ribosomal material was recovered from the lysate using a 35 percent (w/v) high salt sucrose cushion prior to purification using a Ni-IDA column. The RNCs were purified further by applying pelleted RNCs to a butyl HP column equilibrated in a high salt buffer (20mM Hepes, 1.5 M NH4SO4, 400mM KOAc, 12mM MgOAc, 2mM BME, 0.1 percent w/v protease inhibitors pH 7.5). RNCs were eluted using a reverse linear ammonium sulfate gradient into a low salt buffer, which lacked ammonium sulfate. RNC integrity was determined using an anti-his western blot against a series of isolated FLN5 standards as described previously 2.

### Sample preparation and Electron Microscopy

Purified RNCs at ~ 200nM were applied to home-made carbon coated holey-carbon supported grids (C-flat R1.2/1.3, Protochips) vitrified using a Vitrobot Mark IV (ThermoFishcerTM). Data for the FLN5-45 and FLN5-47 RNC complexes were automatically collected on a Titan Krios electron microscope (FEI) operating at 300 kV and equipped with K2 Summit direct electron detector (Gatan Inc.) at the eBIC Diamond light source facility (Harwell Campus, Oxfordshire, UK). The defocus range used was −0.6 to −2.0 μm. Data collection was done using EPU (ThermoFishcerTM). For the FLN5-45 and FLN5-47 RNC samples, movies (45 frames per movie) were collected with a total dose of 45 e-/A2. The pixel size was 1.06 Å/pix. on the sample.

### Image-Processing

Relion 2.0 was used for all steps in image processing. For each dataset, movies were aligned using MotionCorr232. CTFFIND4 33 was used to determine defocus values. Particles were semi-automatically from aligned micrographs using RELION ‘Autopick’ programme 32. The number of selected particles for the FLN5-45 and FLN5-47 RNC complexes was ~ 400,000 and ~ 384,000 correspondingly. Selected images were extracted within boxes of 400×400 pixels. The extracted particle images were subjected to two-dimensional (2D) classification. Subset selection of 188,830 particles in FLN5-45 RNC and 144,460 in FLN5-47 RNC datasets were refined that gave rise to refined 3.0 Å and 2.9 Å maps respectively. The cleaned particle sets were then used for analysis of structural heterogeneity by 3D classification in Relion (as shown in Supplementary Fig. 1). We performed extensive 3D classification (Supplementary Fig. 1) to resolve several NC states within the tunnel and particles from the selected 3D classes obtained were refined independently. The maps used for structure refinements were sharpened by applying negative B-factors of up to −100 using Relion 2.0 32. FSC was estimated, using gold standard approach. Resolutions are reported using the 0.143 criterion (Table 3).

**Table 3.** Cryo-EM structure refinement of FLN5-RNCs.

|  | FLN5-45-1 | FLN5-45-2 | FLN5-45-3 | FLN5-47-1 | FLN5-47-2 | FLN5-47-3 |
| --- | --- | --- | --- | --- | --- | --- |
| EMDB | XXXX | XXXX | XXXX | XXXX | XXXX | XXXX |
| PDB | YYYY | YYYY | YYYY | YYYY | YYYY | YYYY |
| Microscope | Titan Krios (300kV) |  |  |  |  |  |
| Pixel size (Å/pix.) | 1.06 | 1.06 | 1.06 | 1.06 | 1.06 | 1.06 |
| Electron Dose (e-/Å2) | 40 |  |  | 40 |  |  |
| Defocus range (µm) | (-0.8 to -2.0) |  |  | (-0.6 to -2.0) |  |  |
| 3D refinement |  |  |  |  |  |  |
| Software | Relion 2.0 |  |  |  |  |  |
| Number of Particles | 5,410 | 44,732 | 7,824 | 5,712 | 30,047 | 20,653 |
| Resolution (Å, 0.143 criterion) | 3.8 | 3.1 | 3.6 | 3.4 | 2.9 | 3 |
| Resolution (Å, 0.5 criterion) | 6.6 | 3.7 | 5.5 | 6.3 | 3.5 | 3.8 |
| Sharpening B factor (Å2) | -53 | -59 | -49 | -63 | -48 | -42 |
| Model Refinement & Validation |  |  |  |  |  |  |
| Refinement software | Phenix & Coot |  |  |  |  |  |
| All-atom Clashscore | 8.89 | 4.72 | 7.35 | 4.89 | 6.15 | 3.9 |
| Ramachandran |  |  |  |  |  |  |
| Outliers | 0.23 | 0.37 | 0.4 | 0.63 | 0.29 | 0.31 |
| Favoured % | 87.9 | 90.22 | 87.91 | 90.85 | 87.08 | 91.02 |
| Allowed % | 11.8 | 9.41 | 11.69 | 8.52 | 12.64 | 8.66 |

### Atomic Modelling

The high-resolution cryo-EM structure of the 70S-p-tRNA-Gly complex (PDB 5NP6)12 was used as a starting model for ribosome structure assessments. The PDB model was rigid body fit into sharpened cryo-EM maps in UCSF Chimera 34 and subsequently refined iteratively using Phenix 35 real space refine and in Coot 36 for improved structure geometry and overall fit. Refined models were assessed using Molprobity scores 37 as well as visual inspection in Coot. For assessment of changes within ribosome tunnel elements, sequence based structure alignments of the fitted ribosome structures were performed in Chimera 34, and RMSD values were assessed to indicate small or large differences. Electrostatic potential maps for ribosome exit tunnel, FLN6 NC and FLN5 structures were calculated using Chimera 34, based on calculating surface maps from pdb models and colouring maps based on charge potentials of each residue. Initial SecM NC model was obtained using pdb model 3JBU 25. The model was rigid-body fit into each cryo-EM map in Chimera and then refined manually (residue by residue) in Coot. The SecM NCs were then extended with six to eight C-terminal residues of FLN6 NC and fitted manually in Coot. The differences in positions of the fitted SecM NC in both the FLN5-45 and FLN5-47 RNCs were assessed by RMSD. The remaining FLN6 NC were traced by extending the NC model with poly-alanine residues, connected to FLN5. Each NC state was traced with poly-alanine chain independently. The NC model was then refined using phenix-real-space refine implemented in Phenix 35. Fitting for FLN5 NC density was done using rigid-body fit in Chimera, using the crystal structure (PDB 1QFH) 23. First, the pdb model was manually fitted to reflect C-N terminus vectorial emergence from the ribosome. Step-wise cross-correlations scores were computed using the ‘Fit in map’ programme at resolutions 10-15 Å, assessing translation and rotation shifts that helped to identify the best fit of model in to map. Distances between the FLN5 domain and the ribosome tunnel to determine relative positions of the folded FLN5 NC were determined using Chimera. Distances were measured between A490 nucleotide from H24 rRNA (tunnel vestibule) and C747 in FLN5. FLN5 and FLN6-SecM NC were subsequently connected as a single NC. Figures were prepared using Chimera (UCSF) 34.

## Supplementary Figures

**Supp.Fig.1.**
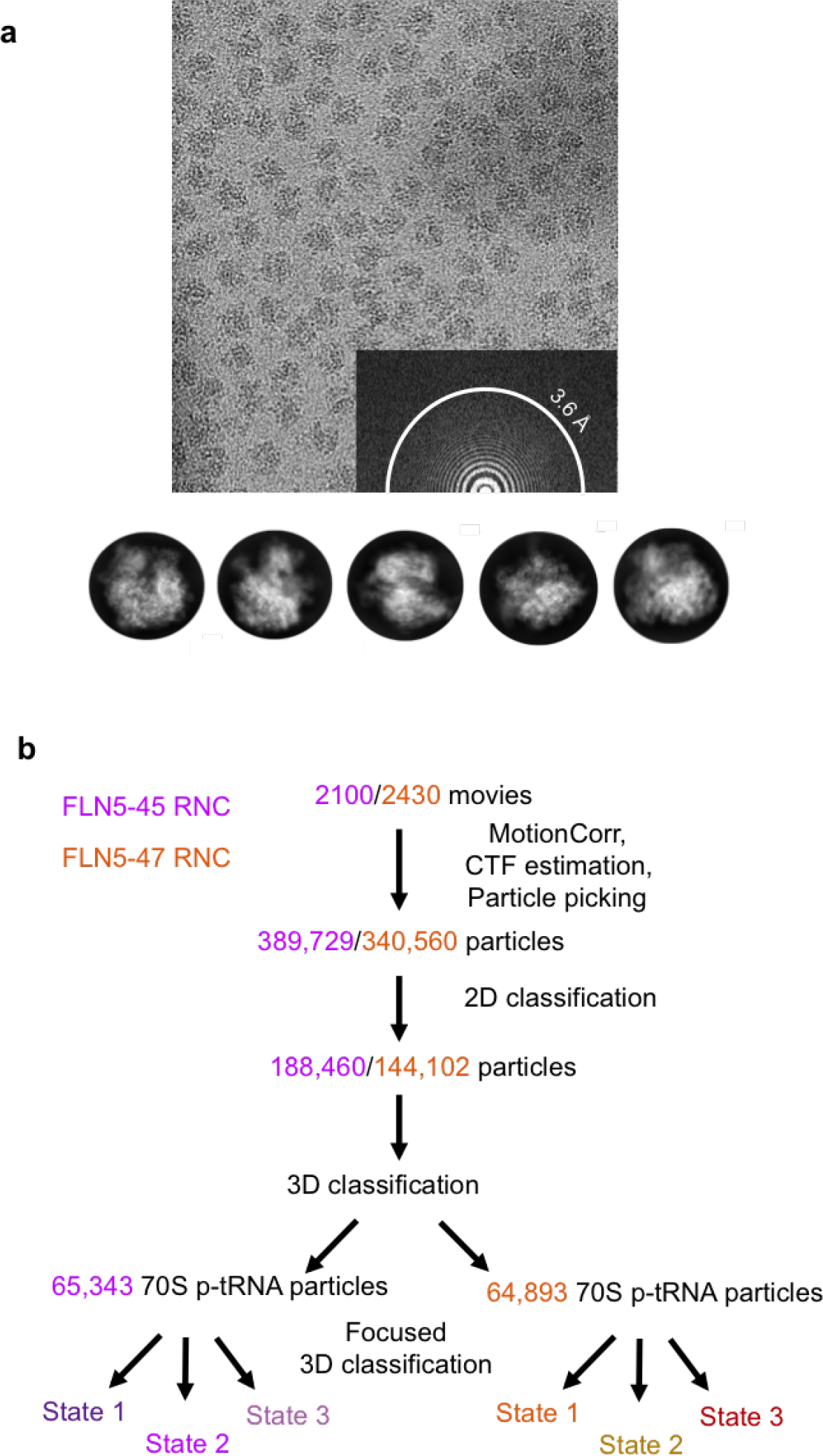
EM analysis of FLN5 RNCs. a. Representative cryo-EM micrograph of FLN5 RNC with its power-spectra, with Thon rings visible at (and beyond) 3.6 Å. Bottom panel shows representative 2D class averages of FLN5-RNCs in classical ribosome orientations, showing high-resolution structural features. b. Image processing scheme for FLN5-45 (magenta) and FLN5-47 (orange) RNCs.

**Supp.Fig.2.**
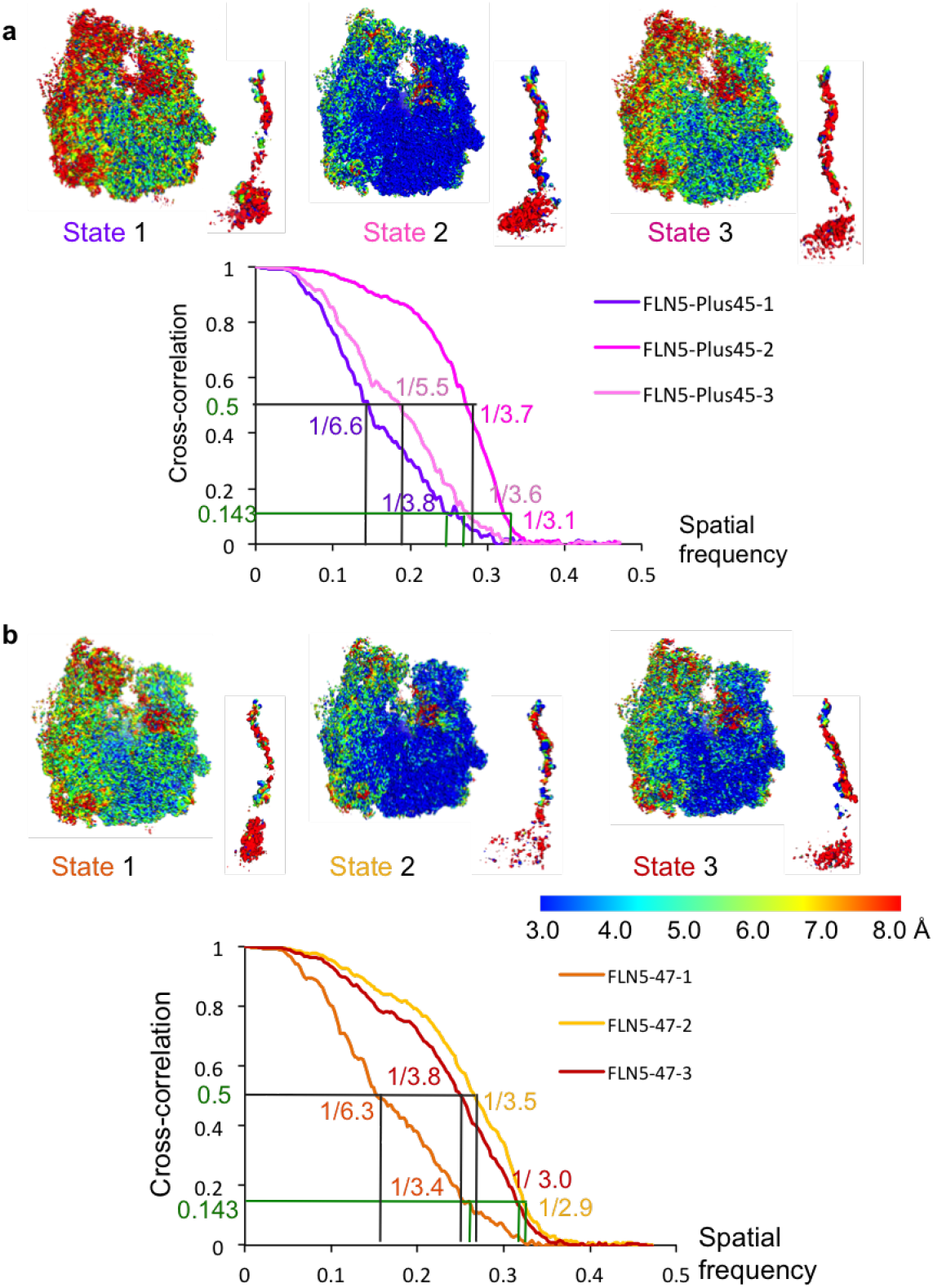
Local resolution and FSC of FLN5-RNCs. Upper panel shows local resolution maps of FLN5-45(in a) and FLN5-47 (in b). RNC states, coloured according to the local resolution across the map. For each RNC state, Fourier Shell-correlation curves are indicated (bottom panels in a and b), with resolutions computed at both 0.5 and 0.143 FSC criterion.

**Supp.Fig.3.**
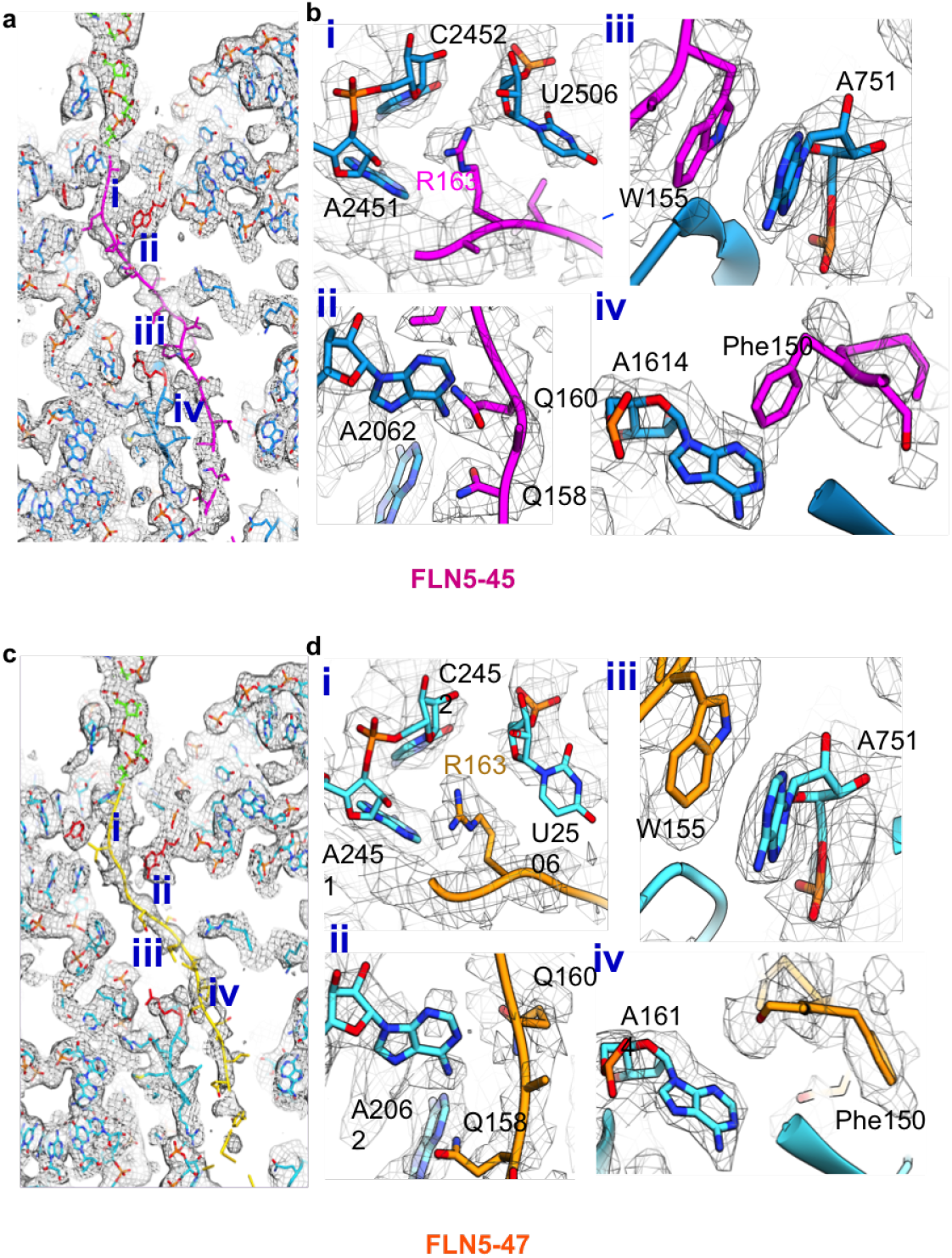
SecM interactions with the ribosome exit tunnel in FLN5-45 and FLN5-47 RNC complex. a, Cross-section of the exit tunnel in FLN5-45 in state 2. b, Interactions of SecM (in magenta) with exit tunnel elements (in blue) are shown in panels i-iv. Map is visualised at sigma value of 1.5. c, Cross-section of the exit tunnel in FLN5-47. Ribosome elements (in cyan) and fitted SecM NC (in yellow) in state 2. Map is visualised at sigma value of 1.5. d, Zoom-in panels i-iv highlight the interaction points between the SecM NC and the ribosome elements lining the upper and central tunnel regions.

**Supp.Fig.4.**
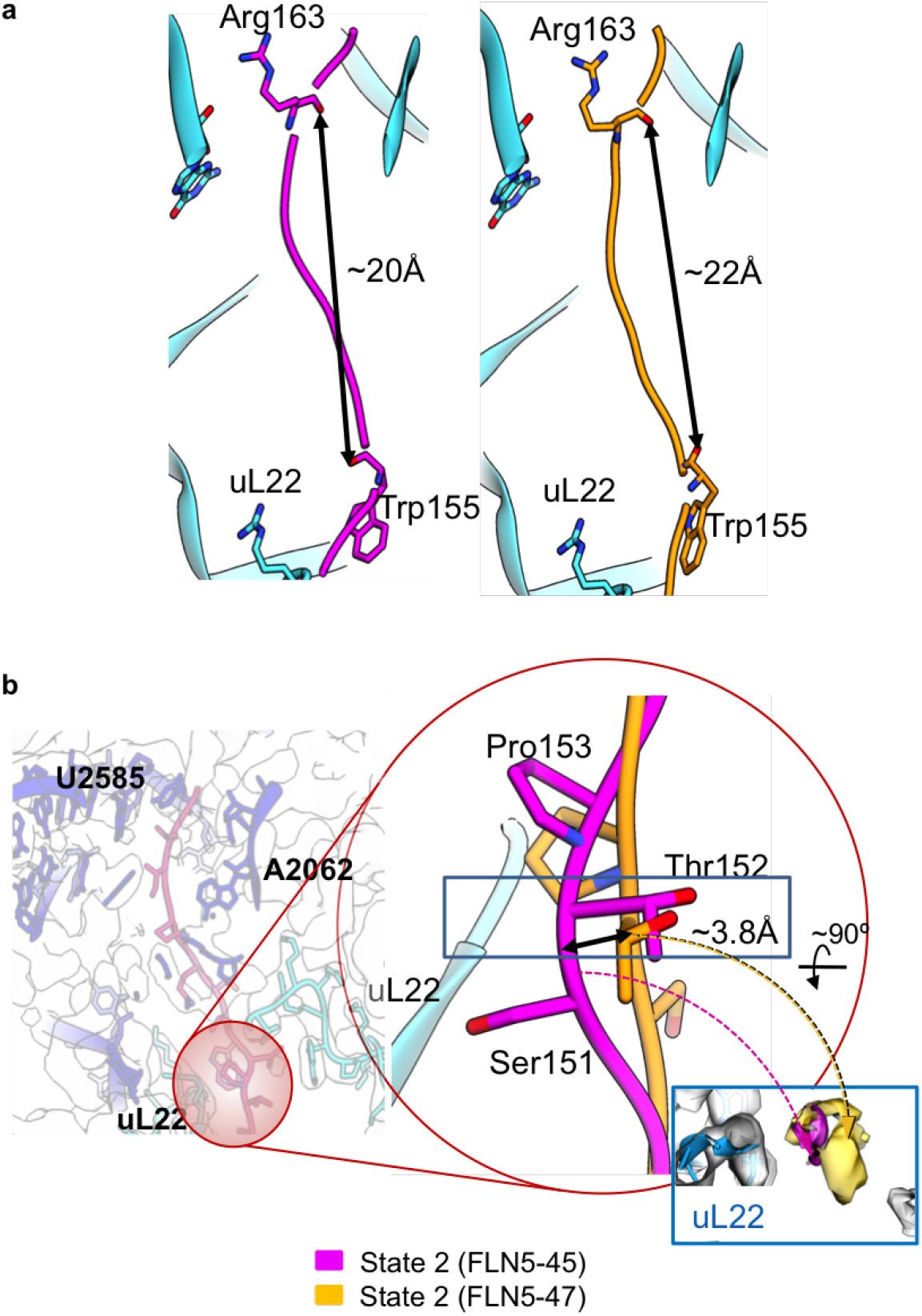
Changes in SecM NC in FLN5 RNC. a, SecM NC states 2 in FLN5-45 (magenta) and FLN5-47 (yellow) RNCs are shown, distances between the 10 residues (from Arg163 to Trp155) indicate NC compaction. Ribosome tunnel elements are colored in cyan. b, Cross-section of the upper tunnel with labelled areas of SecM interactions. A zoom in panel (circled in red) indicates the N-terminal segment of SecM that undergoes a shift of 3.8 Å between FLN5-45 and FLN5-47 RNCs state 2.

**Supp.Fig.5.**
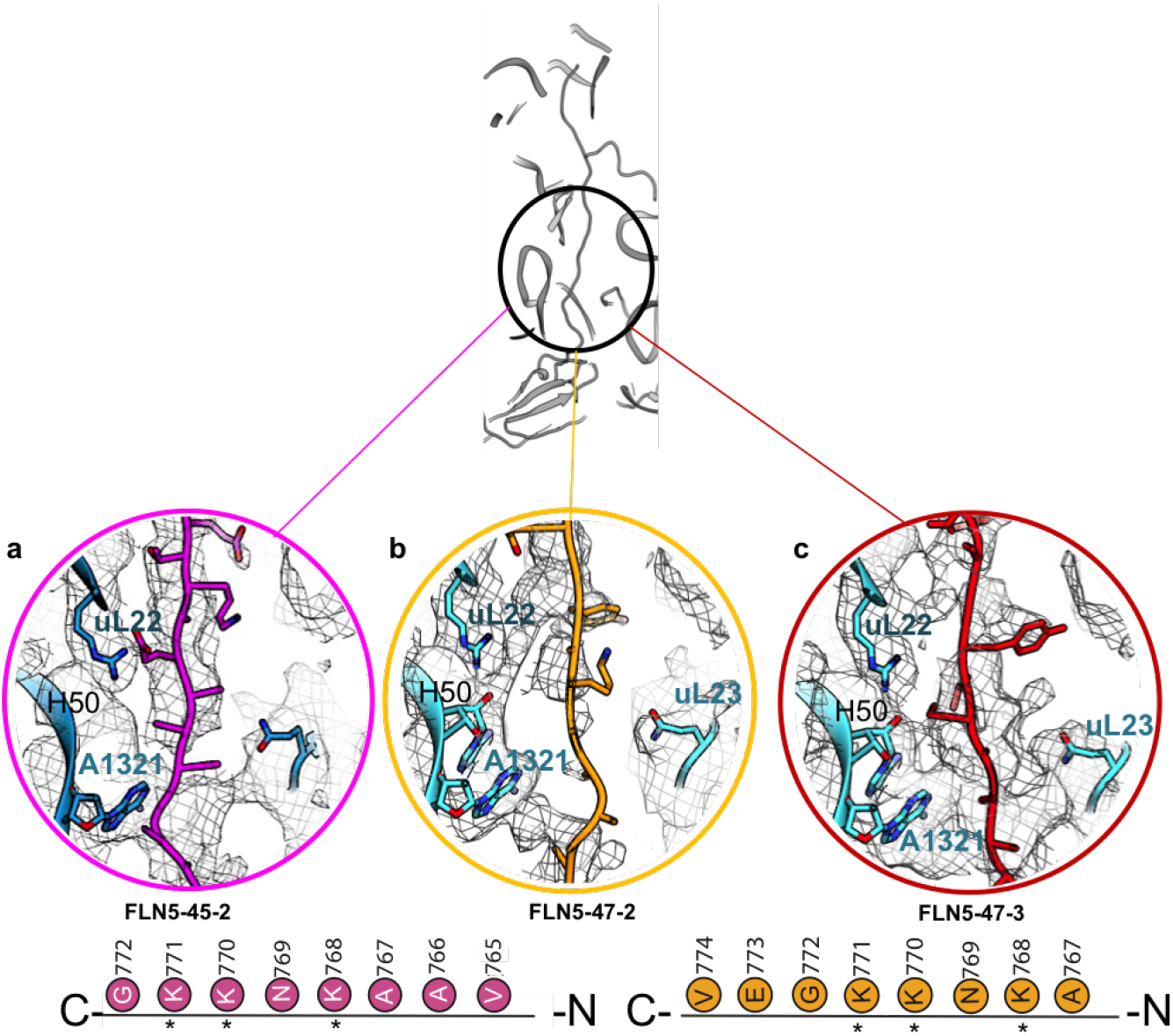
Bifurcation point of FLN6 NC at the ribosome exit tunnel. Top panel shows the region of the ribosome exit tunnel (black circle) occupied by nucleotides of H50 and residues of uL23. Close up panels a-c show EM densities and fitted model of ribosome and NC where interaction points are observed in state 2 of FLN5-45 (in magenta), state 2 of FLN5-47 (yellow) and state 3 of FLN5-47 (red) respectively. Bottom panels show sequences of the FLN6 linker corresponding to the NC shown in panels a-c. Interactions points between ribosome tunnel and FLN6 NC (a-c) are indicated as asterisk.

**Supp.Fig.6.**
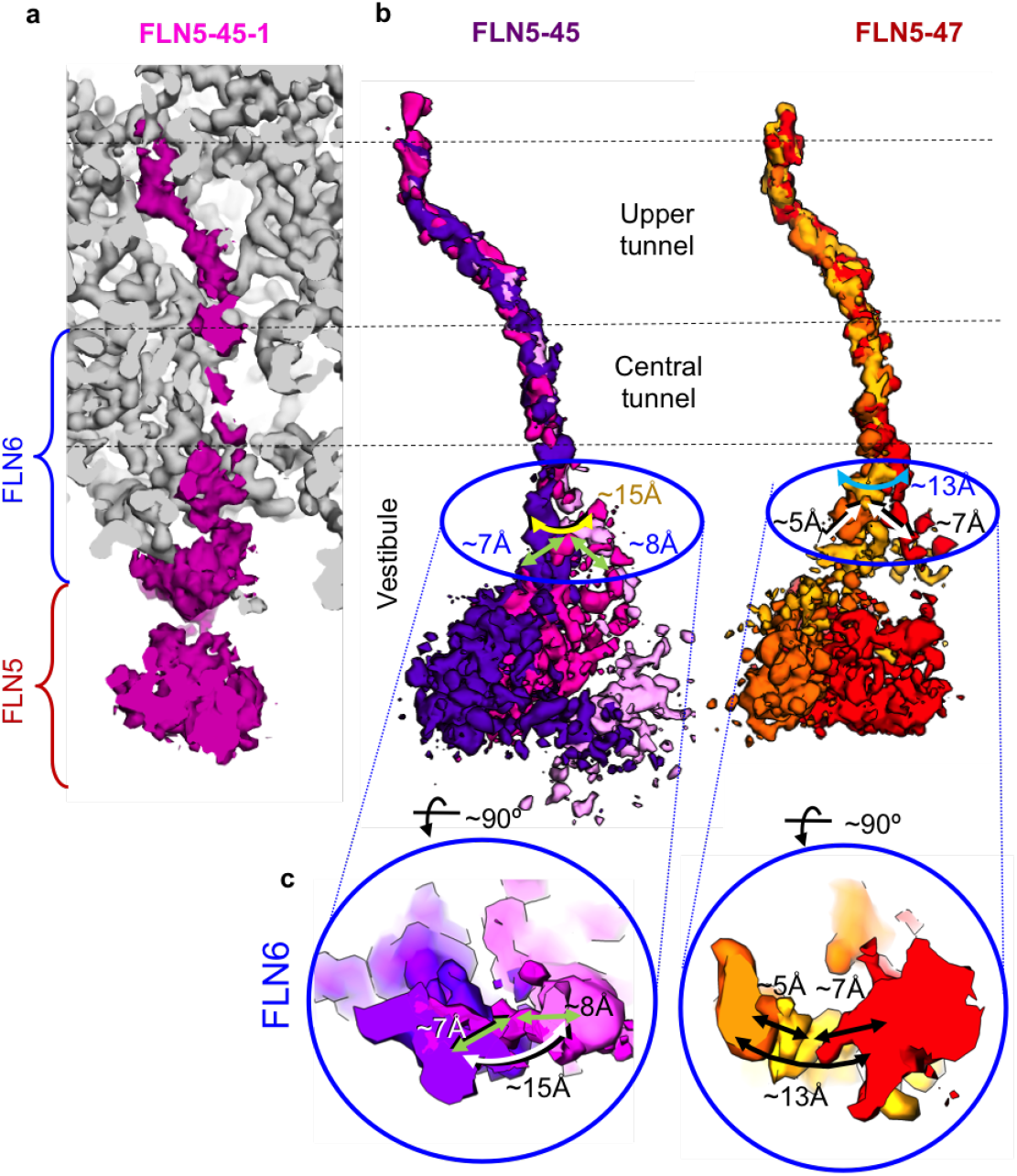
Multiple states of FLN5 NC on the ribosome. a, Cross-section of exit tunnel in the cryo-EM map of FLN5-45 RNC state 1, with ribosome in grey and NC in purple. The map is visualised at sigma value 1, showing FLN6 compaction, attached to a globular FLN5 NC outside the tunnel. b, An overlay of NC maps of states 1-3 in FLN5-45 (left panel) and states 1-3 in FLN5-47 (right panel). Relative shifts between each state is indicated. Shifts between state 1-2 and state 3-2 is indicated by green arrows (FLN5-45) and black arrows (FLN5-47) whilst the shift between state 1 and 3 is indicated by yellow arrow (FLN5-45) and cyan arrow (FLN5-47 RNC). c, Bottom panels show cross sections perpendicular to the axis of the vestibule area (shown in b), with overlay of 3 states, shifts in FLN5-45 and FLN5-47 RNCs are shown.

**Supp.Fig.7.**
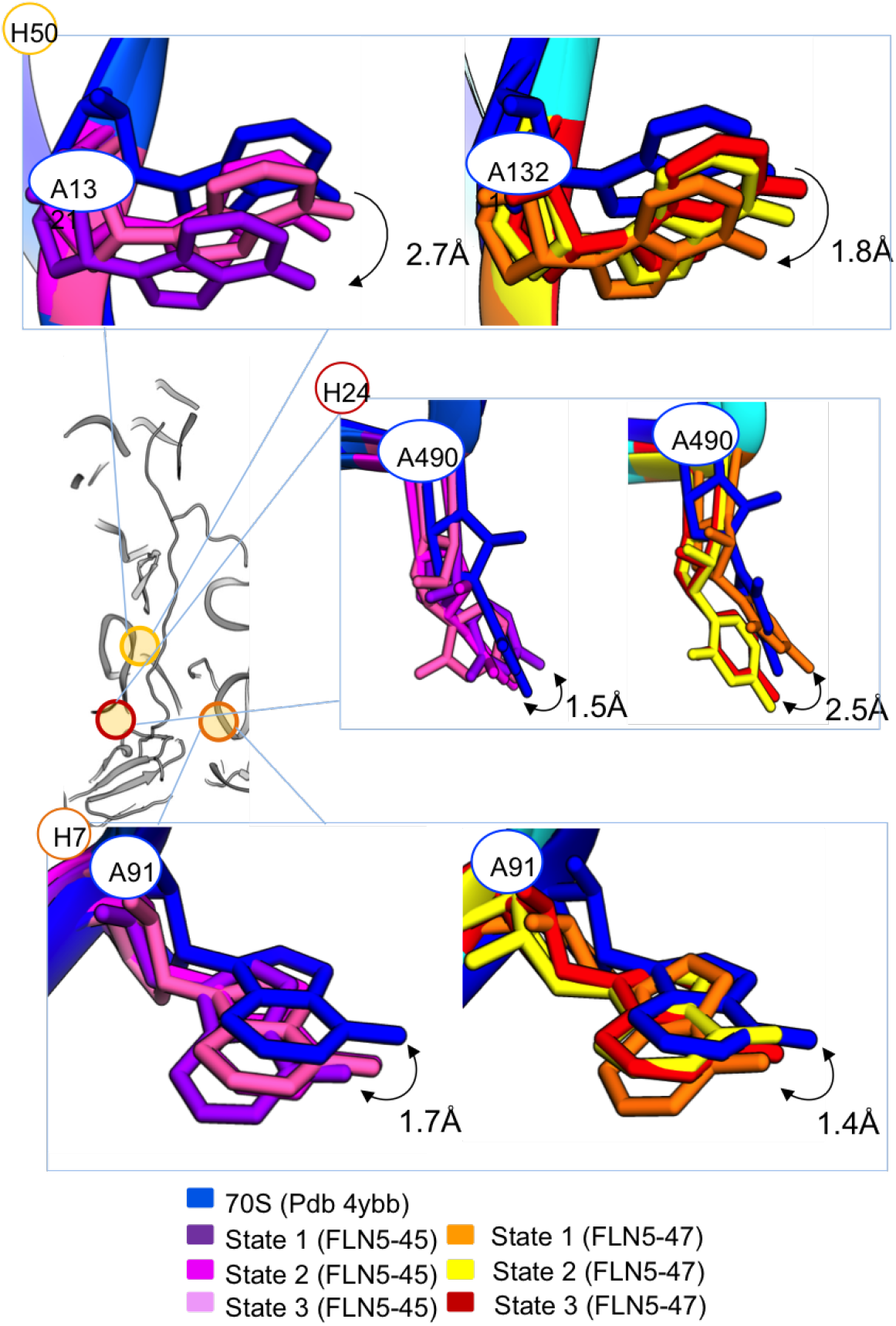
Changes in ribosomal RNA at the tunnel vestibule in FLN5 RNCs. Middle left panel shows exit tunnel cross-section with marked areas. Top panel (H50) shows changes in H50 rRNA at A1321 nucleotide position from empty 70S (blue) and FLN5-45 states (left panel) and FLN5-47 states (right panel). Distances are labelled. Central panel (H24) shows changes in H24 rRNA at A490 nucleotide from empty 70S (blue) and FLN5-45 states (left panel) and FLN5-47 states (right panel). Bottom panel (H7) shows changes in H7 A91 nucleotide from empty 70S (blue) and FLN5-45 states (left panel) and FLN5-47 states (right panel).

**Supp.Fig.8.**
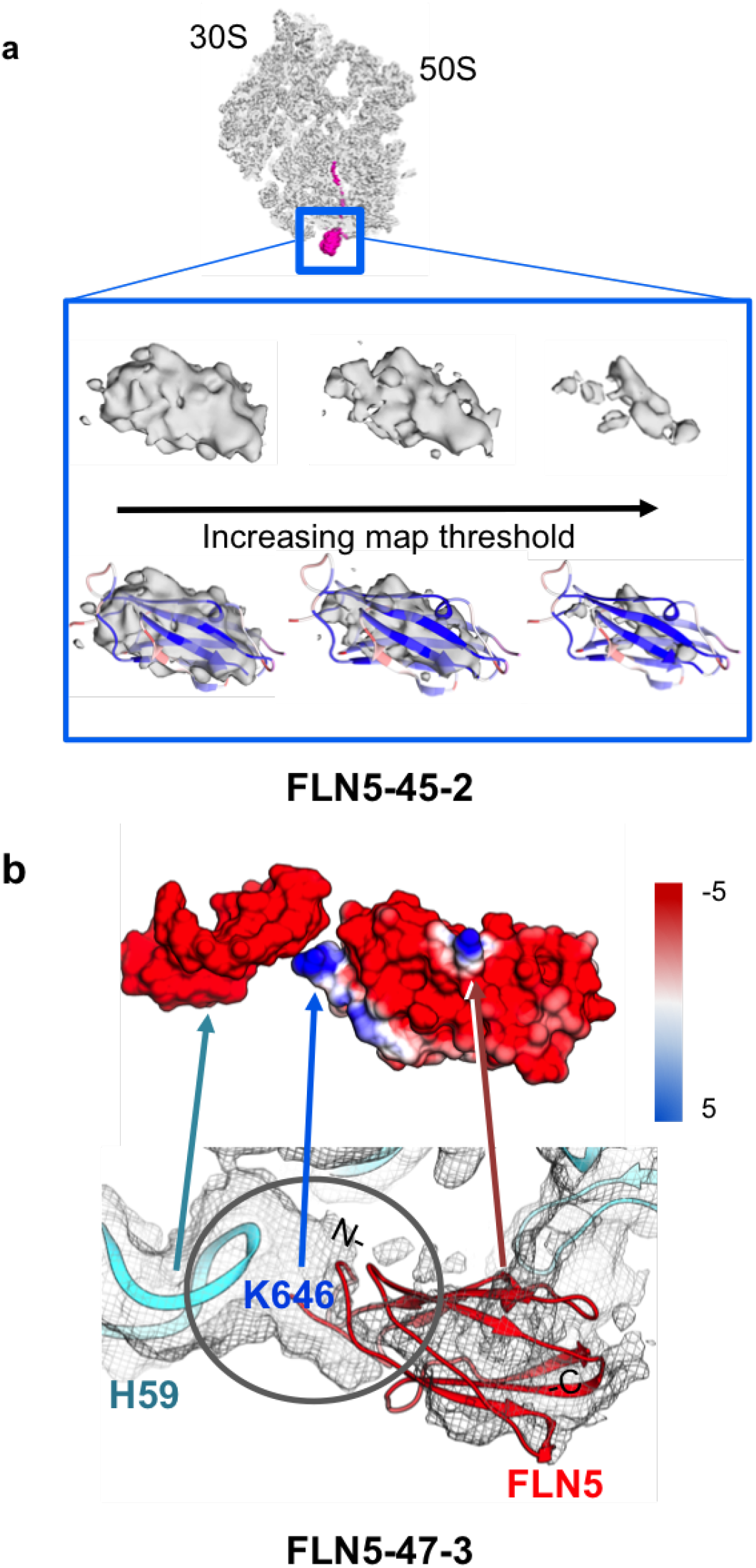
Supplementary Fig. 8. FLN5 close to the exit tunnel. a, Upper panel shows FLN5-45 RNC state 2, ribosome (in grey) with NC density (in magenta). Close-up panel (outlined in blue) shows densities of FLN5 NC (state 2) at three threshold values, with fitted FLN5 model (bottom panel). b, Close-up view into FLN5-47 RNC (state 3) within the area of H59 rRNA, close to exit tunnel. Upper panel shows the electrostatic surface maps for H59 rRNA and FLN5 NC (red – negatively charged, blue – positively charged). Fitted FLN5 domain is shown in red, H59 rRNA domain in cyan. Interaction point between FLN5 and H59 is indicated (grey circle).

